# Plant neighbors differentially alter a focal species’ biotic interactions through changes to resource allocation

**DOI:** 10.1101/2023.11.07.565998

**Authors:** Sophia C. Turner, Jennifer A. Schweitzer

**Affiliations:** Dept. of Ecology & Evolutionary Biology, University of Tennessee, Knoxville TN 37917

**Keywords:** above- and belowground interactions, associational effects, Asteraceae, biotic interactions, competition, interspecific, intraspecific, plant-plant interactions, plant resource allocation, *Solidago*

## Abstract

Plant resource allocation strategies are thought to be largely a consequence of changing abiotic conditions and evolutionary history. However, biotic interactions also influence how a plant allocates resources. As a result, plants mediate indirect interactions between organisms above- and belowground through resource allocation. Neighboring plants can influence plant fitness directly through competition for resources, and indirectly by altering associated community interactions (associational effects). Given the importance of community interactions for plant success, and the known ability for plant neighbors to change these interactions, the goal of this “pandemic project” was to separate inter- and intraspecific plant associations, above- and belowground, to understand how different plant neighbors alter plant resource allocation, and if this in turn alters biotic interactions. We specifically investigated associational effects on herbivory and soil microbial community interactions. To do so, we established a common garden experiment, manipulating plant neighbors and extent of interactions (aboveground only versus above- and belowground interactions, using customized pot types), and measured changes to a focal plant and its biotic interactions over two growing seasons. We found evidence of both neighbor effects and pot type, showing that neighbor interactions affect a focal plant through both above- and belowground processes, and how the focal plant is affected depends on neighbor identity. Though neighbors did not directly alter herbivory or most soil microbial interactions, they did alter the relationship between belowground microbial communities and plant function. Resource allocation responses were reduced with time, showing the importance of extending experiments beyond a single growing season, and is an important consideration when making predictions about plant responses to changing conditions. This study contributes to a growing body of work showing how the community context affects the above- and belowground interactions of a plant through plant resource allocation strategies.

## How we pivoted to a suburban setting to study neighbor interactions

To understand plant neighbor associational effects on Solidago’s above-and belowground interactions, we initially planned a common garden experiment in an old field research site, using seeds from the system. In doing so, we could manipulate neighbor interactions and have access to a realistic source pool of associated communities. Due to travel restrictions imposed, a field experiment was not possible during the pandemic, and so we pivoted to growing this experiment in a suburban backyard. Our study species, *Solidago altissima*, is notorious for its overabundance of interactions with pollinators, herbivores, pathogens and soil microbes. By reducing the experiment to confined conditions in a suburban environment, significantly reduced the insect levels. For instance, we found no evidence of galling, one of Solidago’s quintessential interactions. Another constraint was the availability of space, which meant reducing replication and power in the experiment. During the first year of this experiment, the pandemic was in full swing, and we had no research facilities on campus available for use, so soil samples had to be kept frozen in a conventional household freezer (∼-18 C). Amidst the move of samples to campus, a freezer failure and freezer cleanout, one bag of samples (∼10 samples) disappeared. Fortunately, two years of data compensated for this disappearance. Two important advantages to the backyard experiment were that plant care and watering could be managed on a day-to-day basis instead of regular trips to a field setting, resulting in a very well-attended experiment. Also, we had a dog on the property, who kept the groundhog, squirrel and rabbit activity in the experiment at bay. Although, a squirrel did manage to plant a walnut seed in one of our pots over the first winter, this pot was removed from analysis. Overall, this two-year experiment, despite the constrains of the pandemic, advances our knowledge on neighbor associational effects by showing that the processes that lead to neighbor effects are species specific, largely aboveground rather than belowground, and that effects are reduced over time.

## Introduction

Resource allocation is a critical physiological process that plants undertake to distribute acquired resources, such as carbon and nitrogen, to growth, reproduction, or storage, and is primarily used to optimize fitness (i.e., the ability to successfully reproduce) under different conditions (Bazzaz et al. 1987, Monson et al. 2021, Hartmann et al. 2020). Theory predicts that a plant will allocate resources based on limiting factors (Freschet et al. 2015; Revillini et al. 2016), generally based on abiotic limitations (Tilman, 1982; Gleeson and Tilman, 1992). Therefore, if light limited, a plant would invest carbon in plant growth toward the light, if nutrient limited, a plant would increase fine root production to increase fungal mutualisms, thereby increasing nutrient uptake (Mikkelsen et al. 2008). The response of a plant to limiting factors is both environmentally (i.e., climate and other abiotic factors) and genetically based, with some plants having quick, wide-ranging responses, while other responses can be slower and more restricted (Pierce et al. 2017; Pierce & Cerabolini, 2018). Responses to limiting abiotic conditions result in individual variation in plant performance (e.g., growth, flower success, root production), which in turn can scale up to affect a plant’s interaction with other organisms (Pangesti et al. 2013; Aschehoug et al. 2016).

Plants experience a complex web of direct and indirect interactions, that all occur simultaneously, and all affect how plants invest resources (Agrawal et al. 2006; Kos et al. 2015; Loranger et al. 2013). Insect herbivores can induce the production of volatile compounds, but this comes at a cost, reducing a plant’s ability to allocate resources elsewhere (Erwin et al. 2014; Monsson et al. 2021). These changing resource allocation patterns result in organisms above- and belowground being indirectly connected through their usage of plant resources (Wardle et al. 2004; Mulder et al. 2013; Mougi, 2020; Friman et al. 2021). For example, root herbivores indirectly interact with aboveground herbivores, mediated by plant production of volatile compounds, and this relationship differs based on genotypic plant diversity (Ohgushi, 2008; Wurst et al. 2008). As a result, plant resource allocation is a dynamic compilation of trade-offs used to optimize survival, growth and fitness, the consequences of which affect plant-biotic interactions.

Furthermore, neighboring plants can influence plant fitness directly through resource use, and indirectly by altering associated community interactions (also known as associational effects) (Underwood et al. 2020). Direct plant interactions can result in complementarity effects (i.e., the focal plant performing better in species mixtures than in monocultures), resulting in increased plant performance. The opposite pattern can emerge either from intraspecific interactions inducing facilitation (Zhang & Tielbörger, 2019), or by plant neighbors reducing performance through competition for resources or light (Craine et al. 2013; Sauter et al. 2021). Since plant species allocate and utilize resources differently, different plant neighbors should elicit differing responses in a focal plant. For example, if a plant neighbor results in high shading, a focal species would invest more resources in growth for light. Conversely, if a plant neighbor is a high competitor or has more belowground allocation to roots, the focal plant could increase fine root production to acquire sufficient nutrients. Therefore, plant neighbor identity can create limiting conditions, driving plant resource allocation.

Changes to resource allocation patterns due to neighbor effects can result in different outcomes for associated community interactions. Plant-herbivore interactions are often associated with the density of palatable plant matter (Andersson et al. 2013; Kim et al. 2015), and so if plant neighbors are not palatable to herbivores, then species mixtures would have lower levels of herbivory compared to monocultures. However, if all plants in the mixture have shared herbivore interactions, then species mixtures will experience higher herbivory. These changes to the extent of interactions can alter above- and belowground interactions between organisms – if a plant experiences higher herbivory, it could increase plant stress, thereby changing allocation to fine roots and root symbionts (top-down cascades). In addition, plant neighbors could increase belowground microbial diversity, which could alter the focal plant’s ability to acquire nutrients, thereby reducing plant performance and increasing herbivory (bottom-up cascades). Thus, complex interactions among neighboring plants and diverse above- and belowground communities that associate with a plant, can all be mediated by resource allocation to various plant traits, the results of which can alter indirect linkages among community members, and potentially, overall patterns of biodiversity.

Given the importance of biotic interactions for plant success (Pangesti et al. 2013; Tao & Roode, 2017), and the known ability for plant neighbors to change community interactions (Hamback et al. 2014; Kos et al, 2015, Underwood et al. 2015), the goal of this paper was to understand how neighbor interactions alter plant performance (allocation, function and fitness), the extent to which associated community interactions are changed by plant neighbors, and if changes induced by plant neighbors alter the relationship between the focal plant’s associated community and function. To do so, we established a two-year common garden experiment, with three common old-field Asteraceae species (*Solidago altissima, Achillea millefolium, Silybum marianum*) to test intra- versus interspecific interactions. We grew all plant combinations in two different pot types that separated above- and belowground interactions to test whether neighbors altered a focal plants interaction through above- or belowground processes. Having two years of data allowed us to analyze both years together (with year as a random effect), as well as separately, to determine if changes to allocation patterns are consistent over time. We hypothesized (Fig. 1) that: 1a) the focal plant will elicit a different resource allocation response based on the identity of a plant neighbor, using pot type to determine whether these changes were a result of above- or belowground mechanisms, and 1b) that any effects will be increased in year two due to resource limitation; 2a) in polycultures, we expect open pots to have higher soil microbial richness and diversity due to belowground interaction with plant neighbors, and higher herbivory levels due to a higher specific leaf area (i.e., direct changes to associated community interactions); 2b) changes to plant resource allocation induced by neighbor or pot type will result in changes to the relationship between plant function and its associated community, specifically its belowground soil community and foliar herbivory (i.e., indirect changes to associated community interactions).

**Figure 1.**
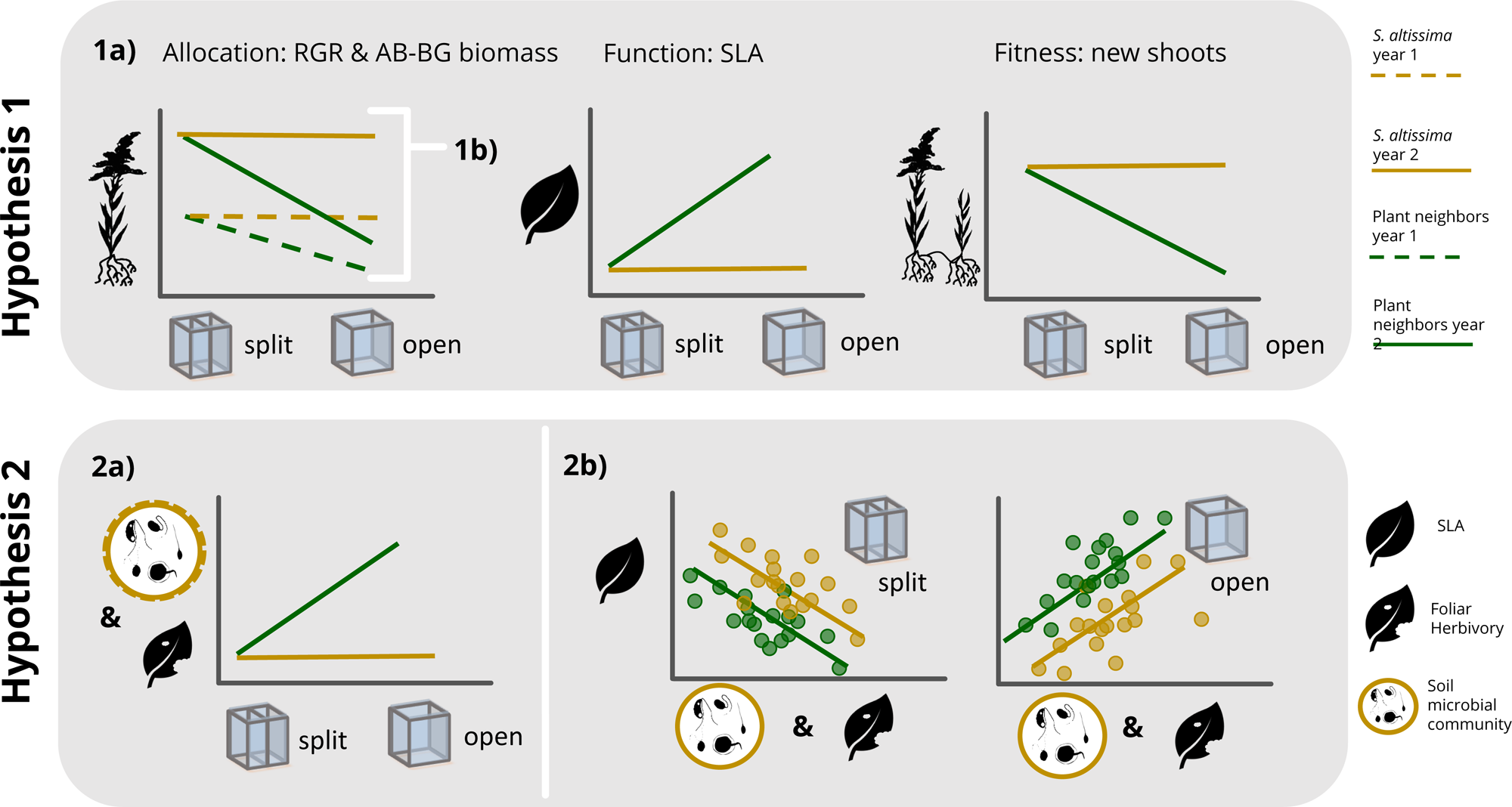
Schematic describing the predictions of our hypotheses, 1a) tests direct changes to plant allocation, function, and fitness, predicting reduction in growth and fitness and increase in SLA in the presence of plant neighbors, mediated by belowground. 1b) predicts that changes to allocation will differ from year to year, with a larger effect in year two as resources become limiting. Hypothesis 2a) predicts an increase in herbivory and soil microbial richness and diversity in open pots with belowground interactions, and hypothesis 2b) predicts that the relationship between the soil microbial community and SLA (plant function) and herbivory (aboveground biotic interactions) will change based on pot type and neighbor. We expect to see an increase in soil microbial richness and diversity in open pots with belowground interactions, and an increase in SLA leading to an increase in herbivory. We expected the opposite pattern in split pots, without belowground interactions with plant neighbors reducing microbial richness and diversity. Yellow lines denote *S. altissima,* green lines denote plant neighbors, and dashed lines refer to year one data.

## Methods

### Study Species

In May 2020 (during the SARS CoV-2 [COVID-19] lock-down) we set up a common garden experiment, with *Solidago altissima* (Tall Goldenrod), a ubiquitous and ecologically important native plant species in Eastern Tennessee, as a focal species. *Solidago altissima* was grown with *Achillea millefolium* (Yarrow), a naturalized species in the region (USDA 2022), as well as a noxious, non-native species, *Silybum marianum* (Milk thistle, USDA 2022) to address specific hypotheses about neighbor interactions. All three species are in the family Asteraceae, are insect pollinated, and grow in similar, disturbed, old-field habitats. *Solidago altissima* is a useful focal species for this study as it is a fast-growing perennial, that has high phenotypic diversity (Etterson et al. 2008). Research has shown the influence of interspecific genotypic diversity (i.e. neighbors) on above- and belowground biomass and other traits of *Solidago* spp. (Genung et al., 2012, 2013), suggesting that *Solidago* patterns of carbon (C) allocation and phenotype may be mediated, in part, from neighboring species.

In this experiment the focal plant, *S. altissima,* was grown from rhizomes, and the plant neighbors, *A. millefolium* and *S. marianum* were grown from seed. Each species was grown in separate, covered trays with a generic potting soil consisting of equal parts peat, vermiculite, and perlite (Premier Pro-Mix BX) and given equal amounts of water. After five weeks, we planted the seedlings into pots in a suburban backyard, using the same Pro-Mix BX, filled to the top of the pots, leaving a lip of 3 cm to prevent any splash over effects between pots, and inoculated with a tablespoon (∼15 g) of a common field soil per plant. Field soil used for the inoculation was collected using an Oakfield soil sampler at a depth of 15 cm, from Forks of the River Wildlife Management Area in Knox Co. Tennessee. Three Ziploc gallon bags of soil were collected in total, and all three were homogenized in a sterile container before application to the experiment. At the time of soil collection, the field was recently mown, and therefore the soil was not associated with a known plant species. The purpose of this inoculum was to represent a common local soil microbial community to see if differing plant resource allocation patterns (as a result of plant neighbor), altered the microbial community through time. The collected soil was refrigerated at 5° C until applied to the experiment (less than 10 days later). In July 2020, we collected *S. altissima* rhizomes, from the same area as the field soil inoculum, but a different section of the preserve. Rhizomes were cut to similar lengths (∼ 10 cm), dipped in rooting hormone (Bonide Bontone II Rooting Powder Plant Growth Regulator), and propagated in Pro-Mix. After four weeks, we added *S. altissima* plants to the potted experiment, also inoculated with the same field soil (stored at 5° C until planting) used for plant neighbors. By planting *S. altissima* four weeks later, we can test how the presence of different plant neighbors affects *S. altissima* establishment and patterns of carbon allocation. We ran the experiment over two growing seasons to determine if observed patterns changed through time.

### Experimental design

The common garden experiment was established in a sectioned off part of the first author’s backyard. The area used was a 10 m x 8. m, level area nearby a water source. Shading from the house and a tree on either side of the experiment occurred, but based on the aspect of the house, shading occurred evenly across the experiment, with some receiving morning shade and others receiving afternoon/evening shade. To account for this, pots were rotated once in each growing season. We found no evidence of shade affecting the data. Pots were placed atop pallets, to be slightly raised from the ground, thereby preventing fine roots from escaping the pot and interacting with a new soil community. Due to space constraints, pots were placed 0.3 m apart. We ensured plants from separate pots did not interact with one another, however, being so close together, meant associated community (i.e., herbivore) effects could be an artefact of placement rather than plant neighbour in the pot. Throughout the entirety of the experiment, plants were watered equally, as needed, with tap water. Watering varied from three times a day in the height of summer year one, to only once a morning, to every few days. We added a slow-release fertilizer to the experiment once, in the summer of year two (Osmocote Smart-Release Plant Food, NPK = 14-14-14, 1 tablespoon per pot).

We used customized, cube-shaped pots in this experiment, made from polypropylene plastic, with a length, width, and height of 0.33 m. Half of the pots had an additional sheet of polypropylene secured in the middle of the pot, dividing the pot in half; silicone adhesive was also applied to all seams to reduce water movement. This divider is both water- and airproof, thereby preventing any belowground interactions between plants on each side (but allowing aboveground interactions; hereafter called “split” pots). These pots were used in previous experiments (e.g., Genung et al. 2013), and the design was shown to not reduce total biomass compared to pots without the divider. The other half of the pots had no divider, allowing above- and belowground interactions, hereafter called “open” pots.

To address how *S. altissima* resource allocation changed based on interactions with each of the neighbor species, and whether the interactions were above- or belowground, we used a manipulative experiment (Fig. 2), planting four seedlings per pot in three combinations: 1) *S. altissima* X *S. altissima*, 2) *S. altissima* X *A. millefolium*, and 3) *S. altissima* X *S. marianum*.

**Figure 2.**
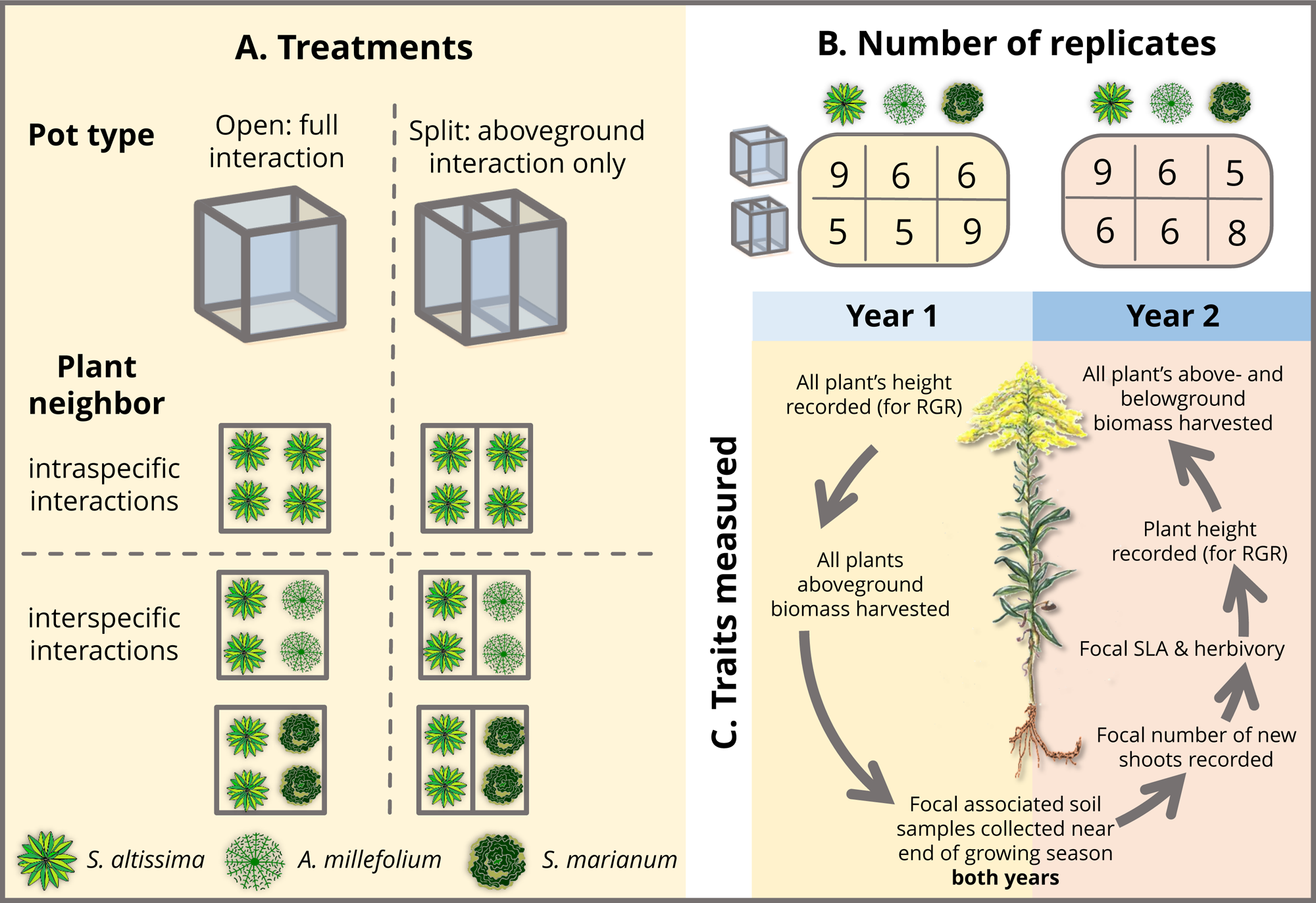
Schematic of the experimental design. Panel A shows the treatments in the experiment - pot type and plant neighbor. We had two pot types, one allowing full belowground interaction (open pots), the other only allowed aboveground interaction (split pots). Our focal species (*S. altissima*) was planted with another *S. altissima* or, *A. millefolium* and *S. marianum,* in both pot types. Panel B shows the number of replicates across treatments and years, as well as the data collected in each year; note, relative growth rate (RGR) was calculated from the height data over time (see methods description). Original *S. altissima* watercolor painting by Shannon Bayliss, adapted for this figure.

The species combinations were planted in split pots (to impede belowground interaction) as well as open pots (to allow interaction). Thus, there were a total of six experimental treatments (3 plant neighbor treatments X 2 pot types). It should be noted that in all pots, *S. altissima* experiences intraspecific interactions, in the polycultures, there are two *S. altissima* individuals with two of the other species. While the monocultures had four *S. altissima* individuals. Our *S. altissima* success rate was low, and as a result we had an uneven sampling design (Fig. 2B), with a total of 5-9 pots per treatment (20-36 plants per treatment). The experiment ran for two growing seasons, for a total of 16 months in the backyard.

### Interpretation of pot type treatment

In polycultures, if the focal plant performance/function is significantly increased in the split versus open pots, this means that plant neighbor reduces the focal plants performance through belowground processes. Conversely, if focal performance decreases in the split pot compared to open pots, this means that the focal plant and its plant neighbor have a positive belowground interaction. If there is no difference in the growth of the focal plant between split and open pots, this means that the plant neighbor does not affect the focal plant’s growth through belowground processes. Furthermore, if the growth in polycultures is lower than in monocultures, then we can surmise that the effect of plant neighbor on the focal plant is through aboveground processes (i.e., competition for light). No significant difference based on plant neighbor or pot type would indicate that the effect of polycultures is no greater or less than that of monocultures.

There are many pathways in which a plant neighbor can alter a focal plant, either directly through resource use or through interactions with other members of the community, or indirectly through associational effects. The mechanisms for these effects can occur either aboveground, such as competition for light or water, or belowground competition for space and resources. Based on our experimental design, we can parse out the likely pathway through which our focal plant is affected by its plant neighbors. Using our study design, we have identified eight possible direct pathways and five indirect pathways of how a plant neighbor could alter focal allocation (Table 1).

**Table 1.** There are many pathways in which a plant neighbor can affect a focal plant and its interactions. Based on our study design, we can interpret how plant neighbor affects plant allocation and function, and/or associated community, and whether it is through above or belowground mechanisms. Using this design, we can detect eight direct and five indirect possible pathways.

| Direct pathways |  |  |  |  |
| --- | --- | --- | --- | --- |
|  | Neighbor effect on Plant growth and morphology | Neighbor effect on associated community | Effect of belowground interaction removal (pot effect) | Theoretical interpretation |
| 1 | no | no | no | Plant neighbor has no effect on a focal plant or its associated community interactions |
| 2 | no | no | yes | Plant neighbors alter a focal plant through belowground mechanisms |
| 3 | yes | no | no | Plant neighbors directly affect a focal plant through resource use, through aboveground mechanisms, but sufficient community diversity exists to result in no effects on associated community interactions |
| 4 | yes | no | yes | Plant neighbors directly affect a focal plant through resource use, without altering associated community interactions, through belowground mechanisms |
| 5 | no | yes | no | Plant neighbors directly alter a focal plants community interactions but there are sufficient resources to result in no direct plant effects (facilitation/resource partitioning), through aboveground mechanisms |
| 6 | no | yes | yes | Plant neighbors directly alter a focal plants community interactions but there are sufficient resources to result in no direct plant effects (facilitation/resource partitioning), through belowground mechanisms |
| 7 | yes | yes | no | Plant neighbors alter a focal plant through direct resource use as well as directly through disruption of associated community interactions, through aboveground mechanisms |
| 8 | yes | yes | yes | Plant neighbors alter a focal plant through direct resource use as well as indirectly through associated community interactions, through belowground mechanisms |
| **Indirect pathways** |  |  |  |  |
| **Scenario** | **Relationship between focal plant and assoc. comm.** | **Neighbor effect** | **Pot type effect** | **Theoretical interpretation** |
| 9 | yes | yes | yes | Plant neighbors alters the relationship between a focal plant and its associated communities, through belowground mechanisms |
| 10 | yes | yes | no | Plant neighbors alters the relationship between a focal plant and its associated communities, through aboveground mechanisms |
| 11 | yes | no | yes | Focal plant interactions with associated community are driven by belowground mechanisms and not effected by plant neighbor |
| 12 | yes | no | no | Focal plant interactions with associated community are driven by aboveground mechanisms and not effected by plant neighbor |
| 13 | no | no | no | There is no distinct relationship between plant and its associated communities, so no effect of neighbor or pot type can be detected |

### Data Collection

We calculated relative growth rate (RGR) on all plants from the first height measurements taken after *S. altissima* was added to the experiment, to the final height data collected before the first harvest (20 weeks). At the end of the first growing season (∼five months after initiating the experiment), we removed the aboveground biomass of all individuals, which was dried at 70° C for 48 h, then weighed. Relative growth rate and aboveground biomass were used as proxies for plant performance. Pots remained uncovered over winter, and weeded as needed. At the start of the second growing season, we counted the number of new shoots that *S. altissima* produced in all pots, using mean shoot number as a proxy for plant fitness. We used height measurements taken at the beginning and end of the second growth season to calculate RGR for year two. At the end of the second growing season, we harvested both above- and belowground biomass. The roots were washed, and all biomass was dried at 70° C for 48 hours and weighed for belowground biomass. These data were used to calculate the root-to-shoot ratio [year two root mass/year two aboveground biomass]). In doing so, we can test whether *S. altissima* altered allocation of resources above- and belowground based on plant neighbor or pot type. During the second growing season, we also collected two random leaves per individual, occurring within the upper quarter of the plant (on the terminal shoot), ensuring they were fully expanded, and representative of the plant. We calculated leaf area using LeafByte (Getman-Pickering et al. 2020), after which, we dried, weighed, and calculated specific leaf area (SLA), an important trait for identifying changes to resource allocation (Firn et al. 2019).

To determine if plant neighbor alters the focal plants above- and belowground interactions (Hyp. 2), we focused specifically on quantifying rates of foliar herbivory and belowground microbial communities in response to neighbor and pot. Herbivory estimates were measured near the end of the second growing season, as a proxy for accumulated herbivory, using the LeafByte app to estimate herbivory based on quantification of leaf area missing (Getman-Pickering et al. 2020). At the end of each year’s growing season, we collected interspace soil between the two target *Solidago* individuals in each pot to a depth of 15 cm with an oat-field sampler. Soil samples were frozen (initially at -18° C, then moved to -80° C) until DNA was extracted for sequencing. By characterizing the belowground community, we can determine if variation in plant growth and belowground resources altered soil microbiome diversity and composition, as well as whether there were changes to indirect relationships between herbivory rate and belowground community diversity.

### DNA extractions and Illumina MiSeq sequencing

All soil samples had DNA extracted using the DNeasy PowerSoil kit, following the manufacturer’s protocol (QIAGEN Inc., Germantown MD, USA). A two-step PCR approach was used for amplicon sequencing. We amplified the V3-V4 gene region of 16S rRNA, using Illumina recommended forward and reverse primers, modified with adapters for the Illumina MiSeq platform (341 F: CCTACGGGGNGGCWGCAG, 785 R: GACTACHVGGGTATCTAATCC, Klindworth et al. 2013) (Eurofins). Amplification success was confirmed by running each sample on a 2% agarose gel (Sigma-Aldrich, St. Louis, MI, USA).

Agencourt Ampure XP magentic beads were used to clean initial PCR products of any unincorporated nucleotides. We then amplified the cleaned products in a second PCR, using the Nextera XT index kit (Illumina Corporation, San Diego, CA, USA). This second-step PCR consisted of 25 ul KAPA HiFi HotStart taq (KAPA Biosystems, Wilmington, MA, USA), 5 ul each of unique combinations of Nextera XT index primers 1 and 2, and 5 ul of initial PCR product, brought up to 50 ul with PCR grade water. Agencourt Ampure XP beads were again used to purify the now indexed PCR products. These products were then quantified on a NanoDrop 1000 spectrophotometer. Amplicons were then pooled for efficiency, quality and quantity, checked on an Agilent Bioanalzyer, and diluted to 4 pM. For each run, the diluted products were combined with PhiX control DNA (Illumina Corporation, San Diego, CA, USA) at a ratio of 20 % PhiX, loaded onto a v3 600-cycle flow cell set for a paired-end read of 275 bases each, then sequenced on the Illumina MiSeq at the University of Tennessee Genomics Core (Knoxville, TN, USA).

### Statistical approach

To address how *S. altissima* resource allocation, function and fitness changed based on interactions with each of the neighbor species, with and without belowground interactions (Hyp. 1, Fig.1), we measured: relative growth rate (RGR) and aboveground biomass in both years, and root:shoot in year two. In the second year, we also measured SLA (function), and number of new shoots produced (as a proxy for fitness).

We first fitted mixed effects models, with the plant traits that were measured in both years (RGR and aboveground biomass) as the response variables, then tested whether they were affected by our predictors – plant neighbor and pot type (fixed effects), with year included as a random effect (Hyp. 1a). RGR residuals were normally distributed, so we used a linear mixed effect model, while aboveground biomass performed better with a generalized linear mixed effect model, using the lme4 package in R (R Core Team, 2020). The Wald’s test was used to identify significant differences in our model (using the Anova function in the *car* package), followed by the Dunnett’s post hoc test for pairwise comparisons.

To address Hyp. 1b we analyzed each year’s data separately. We built multiple regression linear models and used the “Anova” function in the *car* package in R to test for relationships between performance indicators (year one: aboveground biomass and RGR; year two: number of new shoots, above- and belowground biomass, root:shoot, RGR and SLA) and predictors (Fox & Weisberg, 2019). In these models, the predictors used were pot type (open or split), neighbor species identity (*S. altissima*, *S. marianum*, *A. millefolium*), and the interaction between each of these groups. When testing how aboveground biomass and RGR of the focal *S. altissima* changed based on our predictors, we included neighbor aboveground biomass as a covariate in the initial aboveground biomass model, and neighbor RGR as a covariate in the initial RGR model. We included neighbor allocation (aboveground biomass or RGR) to determine if changes to *S. altissima* resource allocation were as a result of the identity of its neighbor or the growth of the neighbor. While the addition of a covariate reduces degrees of freedom, it increased the explanatory power of our models. All predictors were included in the initial model, then the “StepAIC” function in the *MASS* package in R was used to identify the most suitable predictors to keep in the model (Venables & Ripley, 2002). A new model including only the suitable predictors was performed, and an Anova was used to test for significance. In the case where significant interactions occurred, we split the model to identify how interactions changed based on plant neighbor and pot type. Tukey HSD post hoc tests were used to determine significance within groups. It should be noted that *S. altissima* and *A. millefolium* are perennials, and therefore grow new aboveground biomass each year. Whereas *S. marianum* is a monocarpic biennial (sometimes acting as an annual). This was a nutrient poor experiment, to increase the competitive interactions, therefore *S. marianum* acted as an annual. For this reason, pots that contained *S. marianum* in the first year changed to represent only microbial interactions in the open pots (legacy effects of *S. marianum*), and no interaction in the split pots in the second year.

To understand if there were direct effects of plant neighbor on *S. altissima* interactions above- and belowground (Hyp. 2a), we measured foliar herbivory rates in the second growing season using leaves collected for SLA and the Leafbyte app, and the soil microbial community at the end of each growing season (i.e., years 1 and 2) to assess changes in the soil microbiome due to the predictors. We standardized the herbivory estimates by leaf area. Once standardized, the data were normally distributed, and we were able to fit a linear model and use the Anova function to test for the effect of plant neighbor, pot type and their interaction on foliar herbivory.

The soil microbial community from each sample was assessed using the DADA2 pipeline to trim and merge microbial sequence reads, to run quality control checks on data, and group data into exact sequence variants (amplicon sequence variants or ASVs). In addition, we calculated *S. altissima* associated bacterial and fungal diversity using the Shannon-Wiener Diversity Index. The Shannon-Wiener Index is the most commonly used index in community ecology and provides information on both richness and evenness. We then used Anova to test if bacterial and fungal richness and diversity differed based on plant neighbor, pot type, year, and their interactions. Using a principal component analyses (PCoA) with the Bray-Curtis dissimilarity index, we visualized differenced in bacterial and fungal community composition, then tested whether bacterial and fungal composition changed based on plant neighbor, pot type, and year, using Permanovas.

To understand if the relationship between plant function and associated communities is affected by neighbor or pot type (Hyp. 2b), we fit appropriate models and Anova to test how the relationship between SLA and belowground community diversity and SLA, and herbivory changed as a result of plant neighbor and pot type. No significant relationship would indicate that our data is unable to detect a link between plant function and the associated community or an effect of plant neighbor. A significant relationship would show that plant function is affected by the associated community. If significant functional relationship occurred, we further explored whether there was a relationship with plant growth (biomass and RGR). All analyses were conducted in R Studio (R Core Team, 2020).

## Results

### Effect of plant neighbor on focal plant allocation from year to year, function and fitness

Overall, we found evidence of both neighbor effects and some pot type effects, indicating variation in above- vs. belowground interactions, on *S. altissima* resource allocation. The direction resource allocation changed tended to differ based on plant neighbor, highlighting the importance of incorporating neighbor identity when studying associational effects (Cahill, 2022; Mutz et al. 2022). Aboveground biomass was reduced by plant neighbor in year one (Figure 3), and when considering both years together (Hyp. 1a, Table 2), but in year two alone, there was no longer an effect (Hyp. 1b, Appendix SI: Table 1). On the other hand, RGR was driven by pot type (Hyp. 1a, Table 2). Over time, *Solidago altissima* RGR increased as RGR of the plant neighbor increased, but only without belowground contact, i.e., split pots (Hyp.1b, Appendix SI: Table 1). With the presence of belowground interaction (open pots) this changed, and *S. altissima* had a reduced RGR with belowground neighbor interactions, compared to monocultures (Appendix SI: Table 1) suggesting higher competitive effects. *Solidago altissima* RGR in open pots changed depending on plant neighbor, with a significant neighbor identity x neighbor RGR interaction in both pot types, which by year two only occurred in open pots (Appendix SI: Table 1).

**Figure 3.**
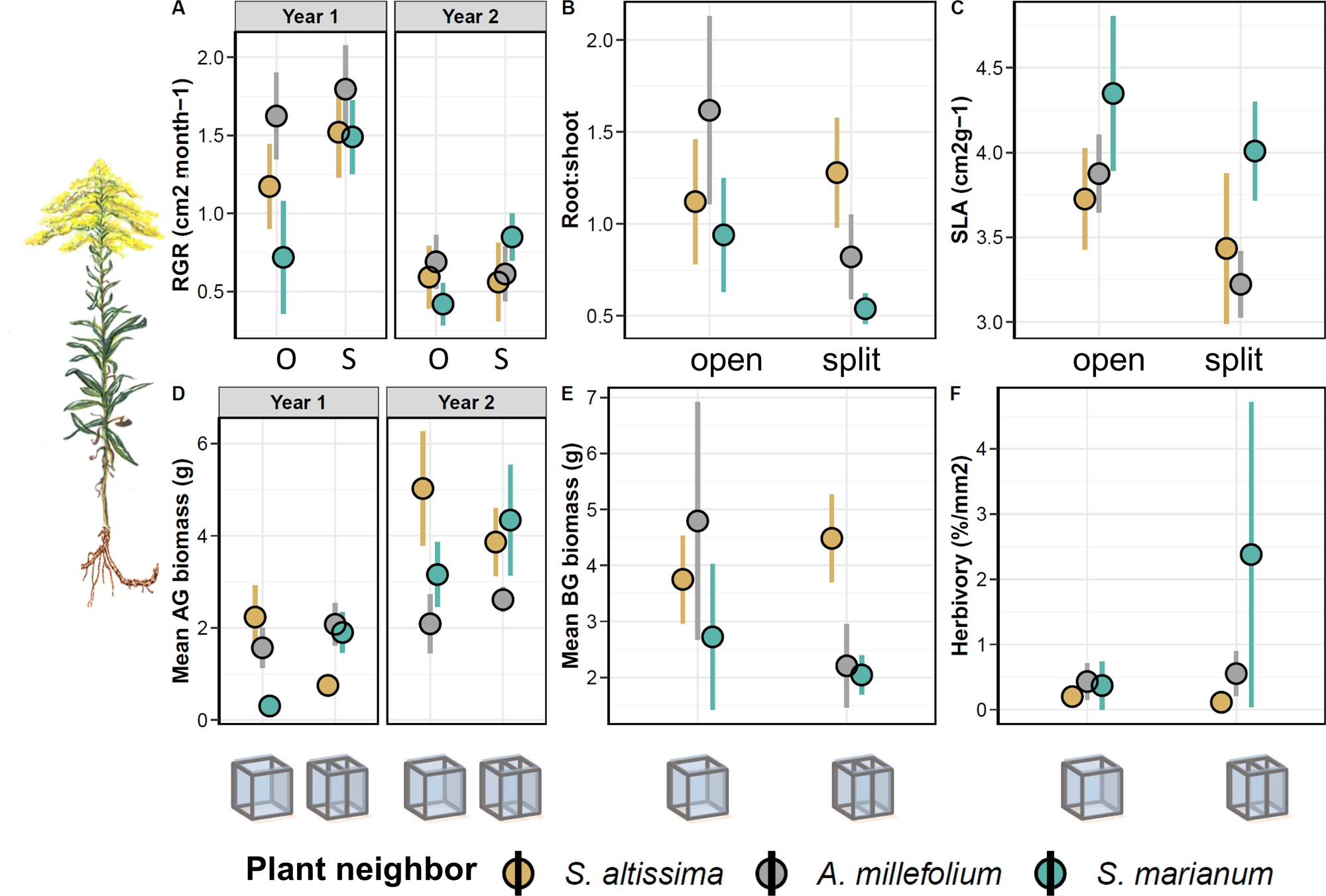
Mean and standard error plots showing *S. altissima* (A) relative growth rate (RGR), (B) root-to-shoot ratio (i.e., root:shoot), (C) specific leaf area (SLA), (D) mean aboveground biomass, (E) mean root biomass, and (F) foliar herbivory as a function of pot type (open or split, with or without belowground interactions, respectively), plant neighbor, (either with *S. altissima, A. millefolium* or *S. marianum,* differentiated by color) and year. Note, all plots show data collected in year two, except for RGR and aboveground biomass which were collected in both years one and two. Original *S. altissima* watercolor painting by Shannon Bayliss, adapted for this figure.

**Figure 4.**
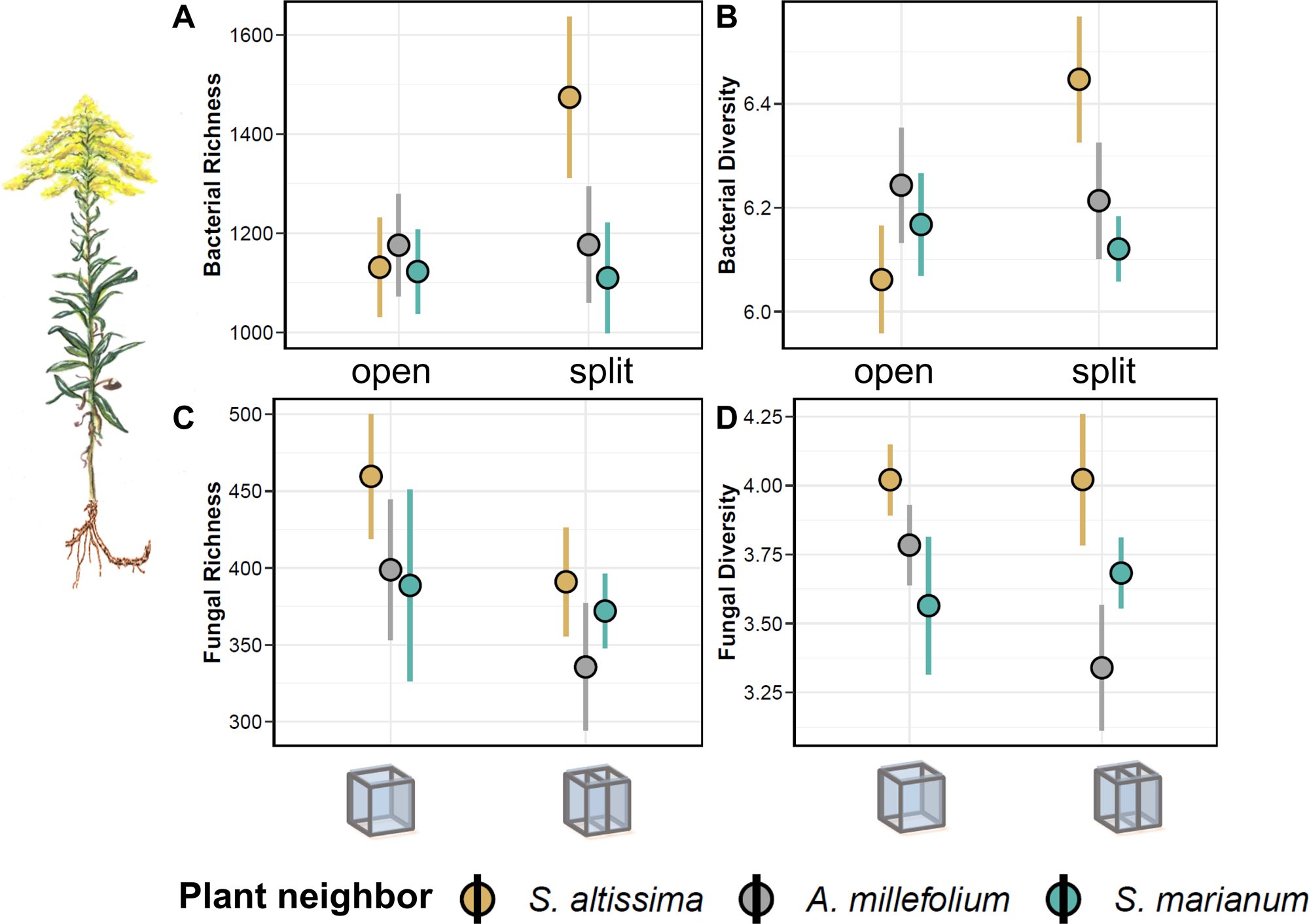
Mean and standard error plots showing bacterial (A) richness and (B) diversity and fungal (C) richness and (D) diversity from soil associated with the focal *S. altissima* based on pot type (i.e., with or without belowground interactions) and plant neighbor; colors represent specific plant neighbors. Original *S. altissima* watercolor painting by Shannon Bayliss, adapted for this figure.

**Figure 5.**
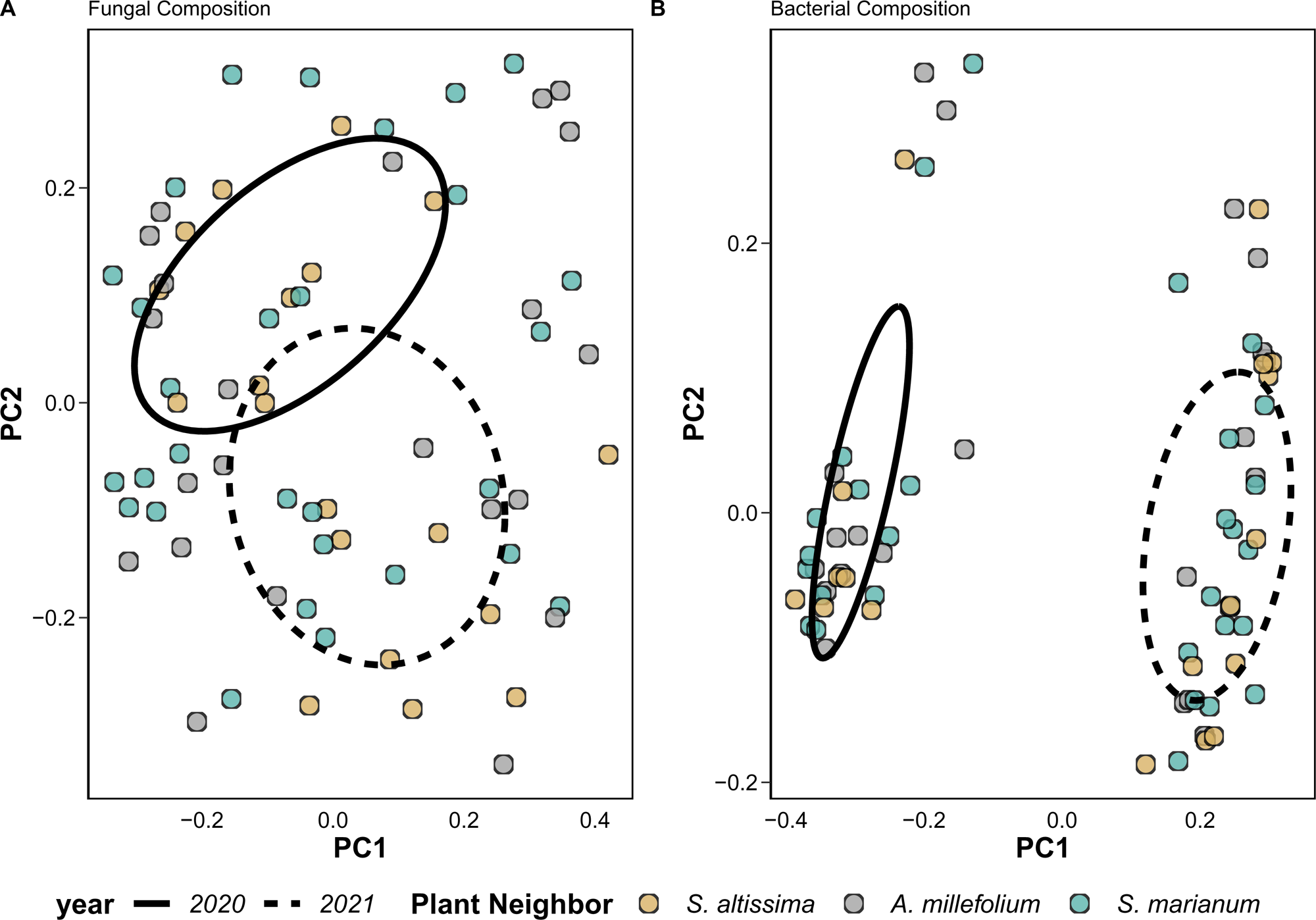
PCoA biplots used to visualize dissimilarities in (A) fungal community composition, and (B) bacterial community composition. Points represent the community composition of each sample (*S. altissima* associated soil from each pot), colored based on plant neighbor (gray for *A. millefolium,* yellow with *S. altissima* and blue with *S. marianum*). The 95% confidence ellipses are used to display the region where 95% of the samples are, for each year. Year was significant for both bacterial and fungal composition, while neighbor and pot type did not alter microbial composition.

**Table 2.** Results from Anovas testing a linear mixed effects model and general linear mixed effect model to predict how the fixed effects of plant neighbor and pot type (i.e., with or without belowground interactions) influence aboveground biomass and relative growth rate (RGR), with year as a random effect in both models. *N*, *P* values are given. Bold types indicate significant effects (*P* < 0.05).

| Response Variables | Predictors | *N* | *df* | *Chi*<sup>2</sup> | *P-value* |
| --- | --- | --- | --- | --- | --- |
| Aboveground Biomass | Pot type | 77 | 1 | 0.194 | 0.660 |
|  | Neighbor |  | 2 | 7.112 | **0.029** |
|  | Neighbor x Pot type |  | 2 | 9.447 | **0.009** |
| RGR | Pot type | 77 | 1 | 0.194 | **0.042** |
|  | Neighbor |  | 2 | 7.112 | 0.230 |
|  | Neighbor x Pot type |  | 2 | 9.447 | 0.225 |

In year two, root biomass, root:shoot and SLA had significant differences in the split pot treatment between *S. altissima* monocultures versus *S. altissima* planted with *S. marianum* - performing better in monocultures (i.e., higher belowground biomass and lower SLA; Fig. 3). There were no neighbor effects of A. millefolium on S. altissima root biomass or SLA Hyp. 1a, Table 2). We found evidence of interspecific competition, as *S. altissima* yielded higher biomass above- and belowground when planted with itself compared to the other plant neighbors.

*Solidago altissima* shoot number was significantly affected by the number of shoots its neighbor had in open pots but not in split pots (i.e., when exposed to belowground interaction, *S. altissima* was negatively affected by the number of shoots that *A. millefolium* produced (Chi^2^ = 4.5124, df = 1, P = 0.034)). While significant, this was a weak negative relationship, with a coefficient of -0.053. Further, *S. altissima* shoot number was significantly higher when planted with itself, or in the legacy *S. marianum* pots (no other plant interaction), compared to pots with *A. millefolium*. Interestingly, when *S. altissima* was grown with itself, irrespective of pot type, it produced significantly more shoots compared to when planted with the other neighbors (Chi^2^ = 5.891, df = 1, P = 0.015).

In most cases, *S. altissima* trait responses were more varied in open compared to split pots, showing that the presence of plant neighbor belowground interactions increased variation in plant response. Taken together, these results support our first hypothesis (1a), that *S. altissima* allocation (above- and belowground biomass and RGR) is reduced with inter- versus intraspecific plant interactions, though the mechanism is not consistent across traits measured or time (1b), and predominately shows neighbor rather than pot (i.e., above- vs. belowground) effects.

### Effect of plant neighbor on foliar herbivory and soil microbial community structure

We found limited support for hypothesis 2a. Herbivory did not differ significantly based on plant neighbor or pot type (Appendix SI: Table 1). Furthermore, both bacterial and fungal richness and diversity did not differ significantly based on plant neighbor or pot type (Appendix SI: Table 2). Results from permanova showed that both bacterial and fungal community composition was mostly driven by year (bacteria: F = 15.438, P = 0.001; fungi: F = 4.8368, P = 0.001), rather than plant neighbor (bacteria: F =0.8293, P = 0.828, fungi: F = 1.1331, P = 0.238) or pot type (bacteria: F = 1.2082, P = 0.152; fungi: F =1.407, P = 0.078).

### Linkages between plant phenotypic change and changes to associated community interactions

We found no evidence of a relationship between *S. altissima* SLA and fungal richness or diversity, nor herbivory. However, we did find a negative linear relationship between SLA and bacterial richness and a quadratic relationship with bacterial diversity (Appendix SI: Figure 1). Neither of these patterns could be explained by plant neighbor or pot type. To understand why there was a quadratic relationship between SLA and bacterial diversity, we explored the relationship between bacterial diversity and other traits. We found that when planted with *S. marianum*, there was a negative effect on the relationship between bacterial diversity and *S. altissima* growth and morphology (Table 3). *Achillea millefolium* had no detectable associational effects on *S. altissima,* through changes to the relationship between S*. altissima* and its associated communities.

**Table 3.** Summaries of linear models showing the effect of plant neighbor and pot type on the relationship between plant traits and bacterial diversity, significant relationships are indicated accordingly: *** p < 0.001; ** p < 0.01; * p < 0.05. All continuous predictors are mean-centered and scaled by 1 standard deviation.

| Model | AG biomass yr 1 | AG biomass yr 2 | RGR yr 1 | Root biomass | SLA | Herbivory |
| --- | --- | --- | --- | --- | --- | --- |
| (Intercept) | 6.34 ***<br>(0.09) | 6.42 ***<br>(0.1) | 6.33 ***<br>(0.09) | 6.35 ***<br>(0.09) | 6.35 ***<br>(0.09) | 6.34 ***<br>(0.09) |
| Pot type: split | 0.06<br>(0.08) | 0.05<br>(0.08) | 0.06<br>(0.09) | 0.08<br>(0.09) | 0.05<br>(0.08) | 0.06<br>(0.09) |
| neighbor: *S. altissima* | -0.02<br>(0.11) | -0.15<br>(0.13) | -0.03<br>(0.11) | -0.07<br>(0.13) | -0.07<br>(0.11) | -0.03<br>(0.11) |
| neighbor: *S. marianum* | -0.24 *<br>(0.1) | -0.34 **<br>(0.11) | -0.23 *<br>(0.1) | -0.26 *<br>(0.11) | -0.23 *<br>(0.1) | -0.25 *<br>(0.1) |
| AG biomass yr 1 | 0.07<br>(0.04) |  |  |  |  |  |
| AG biomass yr 2 |  | 0.08<br>(0.05) |  |  |  |  |
| RGR yr 1 |  |  | 0.04<br>(0.04) |  |  |  |
| Root biomass |  |  |  | 0.03<br>(0.05) |  |  |
| SLA |  |  |  |  | -0.06<br>(0.05) |  |
| Herbivory |  |  |  |  |  | 0.02<br>(0.04) |
| N | 38 | 38 | 39 | 38 | 39 | 39 |
| R<sup>2</sup> | 0.26 | 0.26 | 0.21 | 0.19 | 0.23 | 0.2 |

## Discussion

The overall objective of this pandemic pivot project was to test the effects of inter- and intraspecific plant neighbors on resource allocation of a focal plant, and to determine if changes to allocation altered direct or indirect interactions among diverse plant-associated communities above- and belowground. Plant neighbor effects have long been studied in a variety of contexts (Goldberg, 1987, Tremmel, D. C., & Bazzaz,1993, Yang et al. 2013, Kos et al. 2015, Gough 2006). While many show that plant neighbors can alter a focal plant’s traits and fitness, not all extend to biotic interactions, or if they do it is pairwise interactions (Hausmann & Hawkes, 2009). We found one study that compared effects of plant neighbors on a focal plant’s the above- and belowground interactions (Kos et al. 2015), though it is difficult to compare as this experiment only lasted for three months. Here, we demonstrate how plant neighbors affect a focal plant and some of its biotic interactions, directly and indirectly, and how these effects change over two growing seasons.

### Plant-plant interactions – effects on plant resource allocation

We found that plant neighbors do indeed alter *S. altissima* resource allocation patterns, but how these changes manifest differs based on plant neighbor identity. Species-specific responses have been shown by others (Kos et al. 2015, Kim et al. 2015, Mutz et al. 2022), and is likely due to plants adapting to differing resource-consumer relationships with their neighbors (i.e., plant neighbors use resources differently, therefore the focal plant responds differently). For instance, in year one, plant neighbor reduced *S. altissima* aboveground biomass, but by year two, when *S. marianum* was no longer in the experiment, there was no longer a neighbor effect. In contrast to classic plant competition literature, which favor belowground processes as a source of neighbor effects (Tilman 1990), we found more support for aboveground neighbor effects, for instance, aboveground biomass was driven by neighbor effects in year one. Furthermore, year two belowground biomass, root:shoot, SLA and number of shoots (our proxy for fitness), increased by neighbor but not pot type, suggesting that competition is not for nutrients, and instead is driven by aboveground processes. This shows that even in nutrient depauperate environments, such as this experiment, aboveground mechanisms (such as light and space) can drive neighbor effects. Broadly, naturally occurring *Solidago* complex (spp. *canadensis*, *altissima* and *gigantea*) are known to dominate old field systems, with other studies demonstrating light availability as a key mechanism for this dominance (Eckberg et al. 2023). Our work substantiates the importance of light availability in mediating plant interactions with both other plants and associated communities (Borer et al. 2014).

### Effects on biotic interactions above- and belowground

Analyses showed that associated communities (i.e., herbivory and *S. altissima’s* belowground microbial community) were not greatly affected by plant neighbor nor pot type, showing that plant neighbor had little direct effects on these interactions. In the case of herbivory, this surprising result could either be due to the experiment being conducted in an area with low natural herbivores or because the method we used only considered foliar chewing damage. *Solidago altissima* in natural systems is subject to mining, galling and tunneling by a diversity of herbivores, and a method that incorporated these types of damage would have been more appropriate. No significant difference in belowground richness and diversity could be an artefact of low sample size and high variation within treatments, as *S. altissima* has been shown to harbor high intraspecific variation in plant-microbiome interactions (Foster et al. 2022). It is also possible that all three Asteraceae species share soil symbionts, or that changes to *S. altissima’s* microbial interactions occurred within the roots instead of the associated soil (Hannula et al. 2021). The observed changes in belowground composition from year to year are likely a result of *S. altissima* conditioning the soil (Beals et al. 2023).

Our results show most support for scenario 3 and 4 (Table 1) whereby most of our plant allocation measures showed neighbor effects on the focal plant with no effect on associated communities. Some pot effects were present too, showing plant neighbors can have effects through belowground interactions, such as with RGR. The only observed changes to associated community interactions that we did find were indirect. We found indirect effects of the *S. marianum* treatment, whereby *S. marianum* reduced *S. altissima* SLA, thereby altering the relationship between *S. altissima* function and bacterial diversity. There were no pot type effects, suggesting that this was a top-down indirect interaction. Taken together, these intricate interactions show that *A. millefolium* has direct effects on plant morphology and fitness but not function and associational effects. In contrast, the invasive *S. marianum*, and its legacy effects, showed negative effects on plant morphology, function and indirect interactions. These results show that different plant neighbors elicit differing responses, affecting a focal plant through multiple pathways. While these indirect interactions may seem cryptic, they are not without importance, as it is through a complex combination of many indirect pathways that plant’s function and performance is governed.

### Contextual considerations

Throughout this experiment, we uncovered demonstrable evidence of neighbor effects on resource allocation and the mechanisms thereof, however there are some important considerations when interpreting these results. We conducted this experiment during the height of the COVID-19, in a backyard, while research facilities were not accessible. As a result, we are limited in the inference we can draw from these data due to the context within which this experiment occurred. Two main limitations occurred. We were space-limited in the backyard, leading to lower replication, therefore analyses of each year separately (Hyp. 1b) have lower power. Since this common garden was established in a backyard, there is a significantly lower associated species pool than if the experiment occurred in a more natural setting. As a result, common *S. altissima* interactions were not observed, such as galling, and the presence of goldenrod beetles. This reduction to the type of herbivory could explain the patterns observed in our experiment (i.e., so significant change to herbivory across treatments due to lower levels of herbivory throughout the system).

### Conclusions and future directions

Understanding the role of neighbor interactions in shaping above- and belowground interactions is becoming increasingly important in an era of global change because communities are changing in multiple ways – such as changes to plant communities from species extinctions and introductions, to alterations in plant associated arthropod and microbial communities due to changing climate and mismatches in population dynamics. Neighbor associational effects are commonplace, however, the ways in which we study them are varied. At this stage, most studies, much like ours, show individual focal plant responses to very specific interactions. However, small differences in experimental designs can lead to varied outcomes in the strength and direction of outcomes. For instance, the type of plant-soil feedback experiment (field versus greenhouse) effects plant growth response (Beals et al. 2020), as do soil inoculum preparation (Foster et al. 2022, van de Voorde et al. 2012), making it difficult to move the field toward any predictive patterns. As a result, even experimental studies using the same species are not necessarily directly comparable. To progress above- and belowground ecology, we need to design experiments in comparable ways, measure the same traits at the same stage, and measure biotic interactions in analogous ways over longer time periods. For example, it is not uncommon for greenhouse experiments to run for a single growing season, or a conditioning phase followed by a shorter-term experiment. While single-season studies are important for understanding initial effects of treatments, resource allocation of perennial and biennial plants is a dynamic process and over the course of two growing seasons, plant response, as we showed here, can change quite drastically. For this reason, results from single growing seasons should be taken with caution, and long-term studies need to be implemented to better understand the relationship plant neighbor effects and plant-soil feedbacks (Bardgett et al. 2005; Beckman et al. 2022). Research has shown that traits change with ontogeny, therefore, we can expect biotic interactions to also change as a plant develops (Garbowski et al. 2021). Therefore, to fully understand the nuances of neighbor associational effects, data collection throughout a plant’s development are needed, across trophic levels.

Lastly, this backyard study has shown the importance of plant-mediated indirect interactions among diverse taxa. Similar to studies such as Bezemer et al. (2005) and Pangesti et al. (2013), here we show indirect interactions among diverse groups of taxa are mediated by the resource allocation of the focal plant species. Further, we illustrate how both above- and belowground processes can shape these interactions. These results have important implications for our understanding of the mechanisms through which plant resource allocation governs above- and belowground interactions (Van Dam et al. 2011). In an era of ever-increasing change, gaining a predictive understanding about how plants mediate interactions between seemingly disparate groups of species through carbon allocation is crucial to understanding drivers of community interactions.

## Acknowledgements

We would like to thank the Dennis Breedlove fund for providing financial support for the initiation of this experiment, and the University of Tennessee Student Faculty Research Award for 16S and ITS amplicon sequencing funds. We are grateful to Veronica Brown from the UTK Genomics Core for assistance with sequencing, Stephanie Kivlin, Laura Russo, Christopher Schadt, Dan Simberloff and Shannon Bayliss for feedback on earlier stages of the manuscript, and Renley Roo-Bayliss for using his canine abilities that kept squirrels away from this backyard experiment.

## Author contributions

SCT and JAS conceived the idea for this experiment, SCT established, collected and analyzed data on the experiment, SCT wrote the article with feedback and assistance from JAS.

## Supplementary information

**Figure S1.**
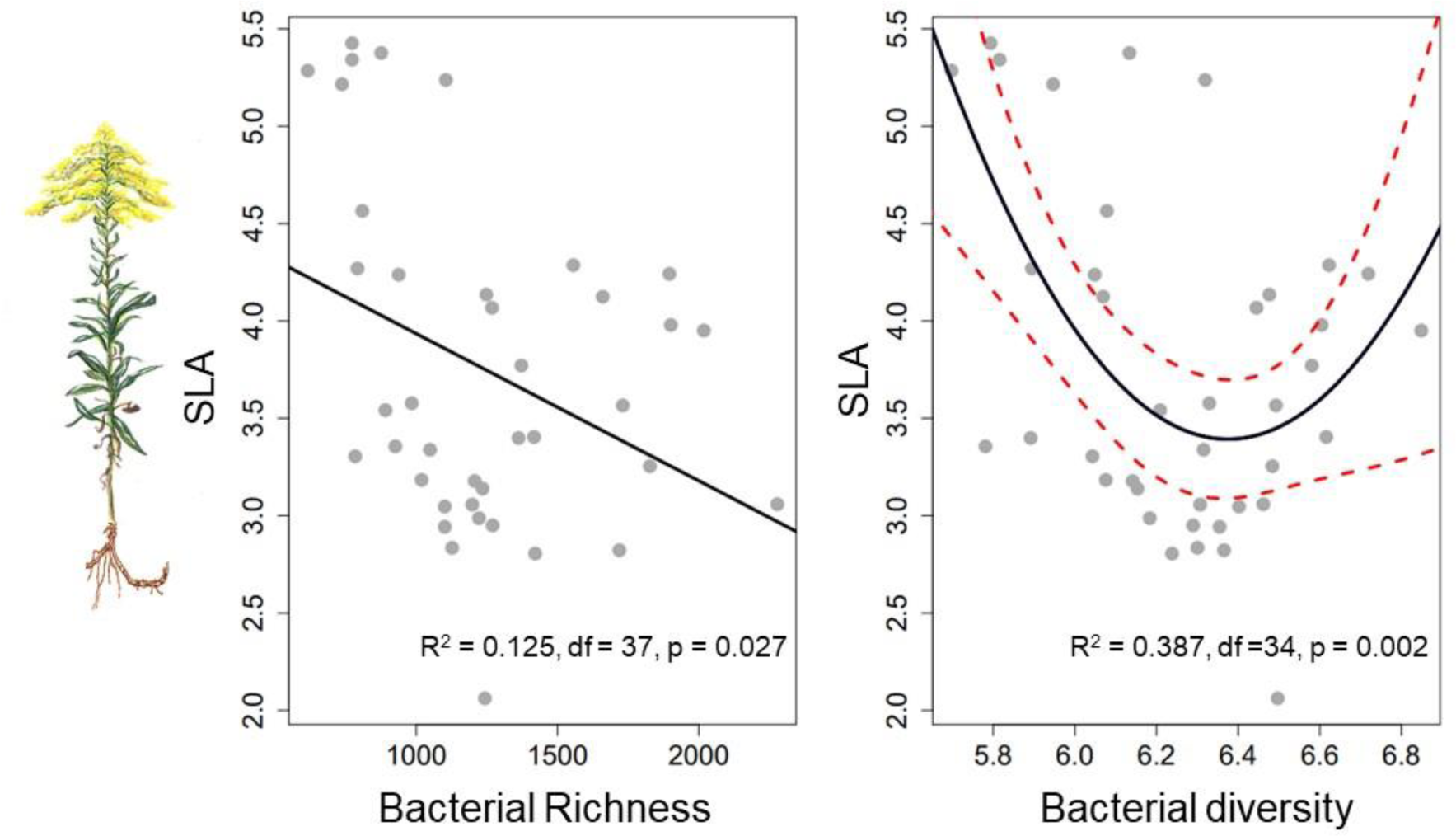
Scatterplots showing the relationship between SLA and bacterial richness and diveristy. Original *S. altissima* watercolor painting by Shannon Bayliss, adapted for this figure.

**Table S1.**
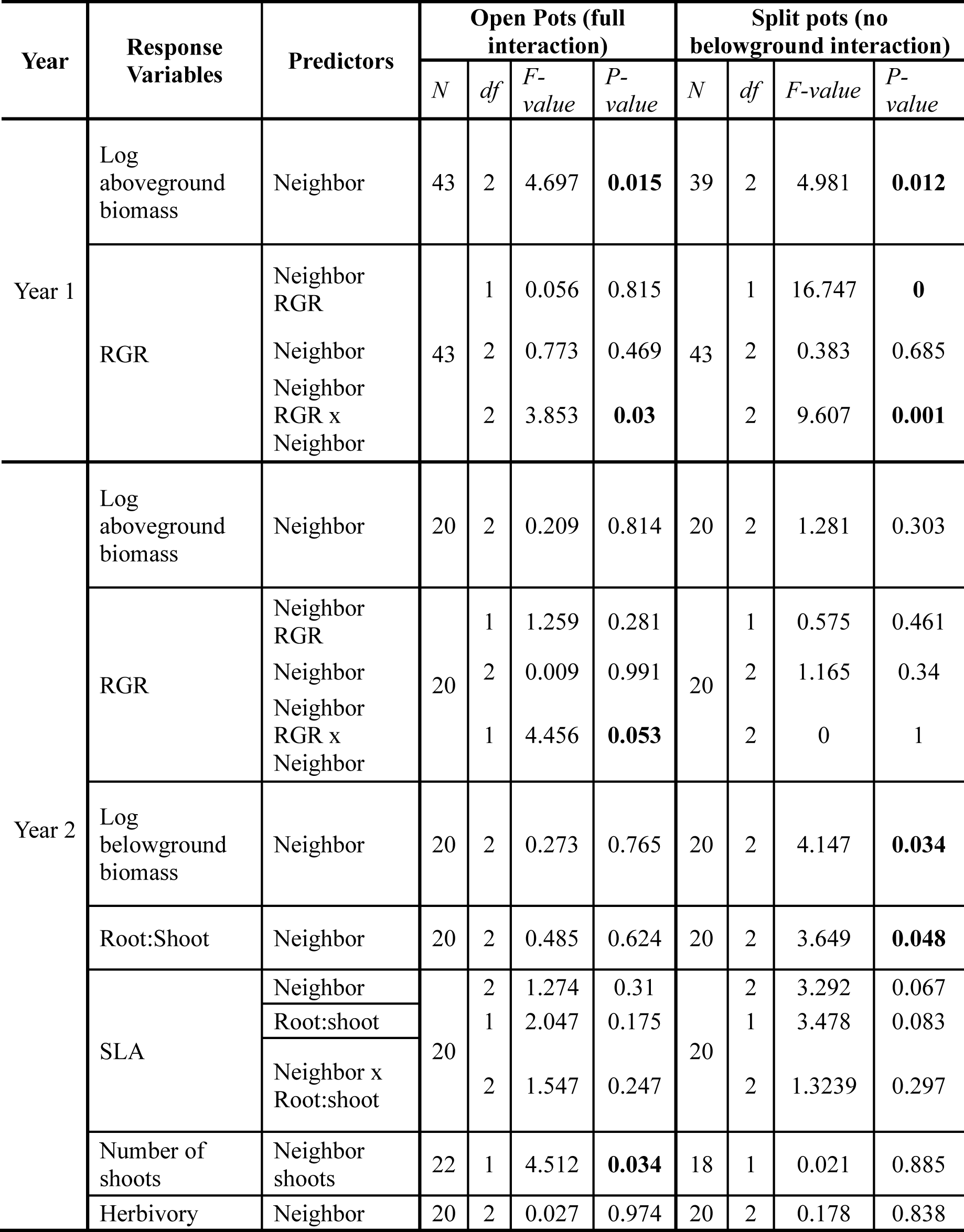
Results from Anovas (car package, R) testing linear models and generalized linear models fit to predict how *S. altissima* plant traits changed as a result of plant neighbor, pot type and their interaction. Bold types indicate significant effects (*P* < 0.05).

| Year | Response Variables | Predictors | Open Pots (full interaction) |  |  |  | Split pots (no belowground interaction) |  |  |  |
| --- | --- | --- | --- | --- | --- | --- | --- | --- | --- | --- |
|  |  |  | *N* | *df* | *F-value* | *P-value* | *N* | *df* | *F-value* | *P-value* |
| Year 1 | Log aboveground biomass | Neighbor | 43 | 2 | 4.697 | **0.015** | 39 | 2 | 4.981 | **0.012** |
|  | RGR | Neighbor RGR |  | 1 | 0.056 | 0.815 |  | 1 | 16.747 | **0** |
|  |  | Neighbor | 43 | 2 | 0.773 | 0.469 | 43 | 2 | 0.383 | 0.685 |
|  |  | Neighbor RGR x Neighbor |  | 2 | 3.853 | **0.03** |  | 2 | 9.607 | **0.001** |
| Year 2 | Log aboveground biomass | Neighbor | 20 | 2 | 0.209 | 0.814 | 20 | 2 | 1.281 | 0.303 |
|  | RGR | Neighbor RGR |  | 1 | 1.259 | 0.281 |  | 1 | 0.575 | 0.461 |
|  |  | Neighbor | 20 | 2 | 0.009 | 0.991 | 20 | 2 | 1.165 | 0.34 |
|  |  | Neighbor RGR x Neighbor |  | 1 | 4.456 | **0.053** |  | 2 | 0 | 1 |
|  | Log belowground biomass | Neighbor | 20 | 2 | 0.273 | 0.765 | 20 | 2 | 4.147 | **0.034** |
|  | Root:Shoot | Neighbor | 20 | 2 | 0.485 | 0.624 | 20 | 2 | 3.649 | **0.048** |
|  | SLA | Neighbor | 20 | 2 | 1.274 | 0.31 | 20 | 2 | 3.292 | 0.067 |
|  |  | Root:shoot |  | 1 | 2.047 | 0.175 |  | 1 | 3.478 | 0.083 |
|  |  | Neighbor x Root:shoot |  | 2 | 1.547 | 0.247 |  | 2 | 1.3239 | 0.297 |
|  | Number of shoots | Neighbor shoots | 22 | 1 | 4.512 | **0.034** | 18 | 1 | 0.021 | 0.885 |
|  | Herbivory | Neighbor | 20 | 2 | 0.027 | 0.974 | 20 | 2 | 0.178 | 0.838 |

**Table S2.** Results from Anovas (car package, R) testing linear mixed effects models fit to predict the effect of pot type (i.e., with or without belowground interactions) and plant neighbor on *S. altissima* associated soil microbial communities (bacterial and fungal richness and diversity), with year as a random effect. Bold typed numbers indicate significant effects (*P* < 0.05).

| Response Variables | Predictors | *N* | *df* | *Chi*<sup>2</sup> | *P-value* |
| --- | --- | --- | --- | --- | --- |
| Bacterial Richness | Pot Type | 69 | 1 | 0.6236 | 0.4297 |
|  | Neighbor |  | 2 | 1.9283 | 0.3813 |
|  | Neighbor x Pot type |  | 2 | 2.5555 | 0.2787 |
| Bacterial Diversity | Pot Type | 69 | 1 | 0.4344 | 0.5099 |
|  | Neighbor |  | 2 | 1.0769 | 0.5837 |
|  | Neighbor x Pot type |  | 2 | 4.4780 | 0.1066 |
| Fungal Richness | Pot Type | 70 | 1 | 1.4440 | 0.2295 |
|  | Neighbor |  | 2 | 1.7031 | 0.4268 |
|  | Neighbor x Pot type |  | 2 | 0.4224 | 0.8096 |
| Fungal Diversity | Pot Type | 70 | 1 | 0.7157 | 0.39757 |
|  | Neighbor |  | 2 | 5.3581 | 0.06863 |
|  | Neighbor x Pot type |  | 2 | 2.4551 | 0.29301 |

